# Bidirectional interconversion between mutually exclusive tumorigenic and drug-tolerant melanoma cell phenotypes

**DOI:** 10.1101/2020.08.26.269126

**Authors:** Yuntian Zhang, Rachel L Belote, Marcus A Urquijo, Maike M. K. Hansen, Miroslav Hejna, Tarek E. Moustafa, Tong Liu, Devin Lange, Fatemeh Vand-Rajabpour, Matthew Chang, Brian K. Lohman, Chris Stubben, Xiaoyang Zhang, Leor S. Weinberger, Matthew W VanBrocklin, Douglas Grossman, Alexander Lex, Rajan Kulkarni, Thomas Zangle, Robert L. Judson-Torres

## Abstract

Human cancers can exhibit phenotype switching, resulting in cells that are more metastatic or that are more tolerant to treatment. However, the relationship between these aggressive states is not well understood. We investigated the dynamics of phenotypic switching in human melanoma cells by monitoring the fluorescent activity from a transgenic reporter of BRN2 promoter activation. Melanoma cells exhibit heterogeneous BRN2 expression patterns that are reestablished upon isolation and clonal outgrowth. Specifically, stable BRN2 expression was generally inherited over multiple generations while undergoing occasional bidirectional interconversion. We found that BRN2 low cells were required for tumor initiation and metastasis in animal engraftment assays but were more sensitive to targeted BRAF inhibition. In contrast, the BRN2 high state was not tumorigenic, but entry into this state was uniform and persistent among cells tolerant to targeted BRAF therapy. Single-cell RNA sequencing analyses revealed core programs exclusive to either the BRN2 high or low cells, each of which is present in *ex vivo* tumors, demonstrating the physiological relevance of these states. Our findings emphasize that one challenge of effectively targeting phenotype switching in melanoma as a therapeutic strategy could be balancing distinct aggressive phenotypes so that sensitizing tumors to BRAF inhibition does not inadvertently lead to further dissemination.

**Teaser:** Unraveling melanoma’s shape-shifting behavior: insights into how cancer cells swap between metastasis and drug evasion.

## Introduction

Phenotype switching refers to a consequential alteration in the transcriptional programs and behaviors of a cell in the absence of additional genetic changes. This process gives rise to tumor heterogeneity and contributes to tumor initiation, primary growth, progression, and metastasis^1,2^. In particular, a variety of phenotypes have been characterized during the progression of cutaneous melanoma - a disproportionally lethal skin cancer arising from neural crest-derived pigmented cells called melanocytes^1,3–15^. Four transcriptional programs frequently associated with melanoma phenotype switching are those controlled by the transcription factors MITF, SOX10 and BRN2 (aliases include POU3F2 and N-OCT3) and the receptor tyrosine kinase AXL^8,16–34^. MITF is considered a master-regulator of the melanocytic lineage and regulates melanocyte differentiation and pigmentation^35^. SOX10 is a well-established activator of MITF expression, but is also critical for the neural crest stem cell (NCSC) program that precedes melanocyte specification during development and plays complex roles in melanoma proliferation and progression^34,36–38^. While the relative expression of the MITF and AXL programs are often described as mutually exclusive^16–19^, their relationship with the BRN2 program is less consistent. BRN2 is a lineage-essential transcription factor that is expressed in melanocyte precursors, silenced in differentiated melanocytes, and heterogeneously reactivated in the dermal component of both benign and malignant melanocytic tumors^20–23^. In both development and disease, BRN2 expression is often, but not always, associated with migration and invasion^24–28^. One model suggests that BRN2 and MITF represent the two poles of a molecular switch that toggles between the distinct states of invasive and proliferative phenotypes^21,27,29^. However, melanoma cells are capable of simultaneous invasion and division^39^. Conditions that induce the invasive phenotype or oppose the proliferative MITF program do not necessitate increased BRN2 expression, nor are BRN2 and AXL obligatorily co-expressed^17,24,26^. In addition, dependent on the context, the genes can also present reciprocal activation or no relationship at all^24,26–32^. These observations collectively challenge the model of a compulsory mutually exclusive MITF/BRN2 axis. Regardless of its relationship to MITF, BRN2 expression oscillates between distinct stages of metastasis *in vivo*, and the gene itself can serve to either promote or suppress melanoma progression^8,23,33^. Thus, dynamic BRN2 expression appears important for melanoma progression, but the contexts in which it serves to promote or restrain aggressive disease remain obscure.

Therapeutic strategies for pharmacological targeting and suppressing specific cellular phenotypes are being explored for combatting melanoma progression^1,14,34,40–44^. However, given that the relationships between distinct and significant phenotypes remain incompletely characterized such strategies could yield unintended, and potentially harmful, consequences. For example, multiple therapeutic resistance phenotypes and multiple tumorigenic stem-like phenotypes have been described^8,16–33,43–45^. Cell surface antigens used to purify tumorigenic cells in some studies prove unreliable in others, and the relationship between drug-tolerant and tumorigenic phenotypes remains incompletely understood^46–54^. We sought to characterize the dynamics of switching between tumorigenic and targeted therapy-tolerant melanoma phenotypes. We reveal a relationship of mutual exclusivity, such that the enrichment of one phenotype necessitates the depletion of the other. Each phenotype was generally inherited within a lineage but could also undergo bidirectional interconversion to re-establish a phenotypic equilibrium within the population. This work reveals potentially significant complications for therapeutic strategies aimed at diverting cells from a drug-tolerant phenotype, as the practice could inadvertently induce a pro-metastatic phenotype.

## Results

### Phenotypic equilibrium is established within clonal human melanoma lines

To monitor the dynamics of phenotypic switching we generated a fluorescent reporter construct for the promoter activity of *BRN2*. We performed ATAC-seq in the human melanoma cell line, 624-Mel, to identify the region of open chromatin upstream of the gene. We defined the proximal active promoter region as spanning from ∼ 817 base pairs (bp) upstream of the transcriptional start site (TSS) to ∼ 195 bp into the untranslated transcript, consistent with prior reports in mouse (**Fig. 1A**)^8^. We cloned this region into a selectable lentiviral mammalian expression construct, adjacent to a nuclear localization signal (NLS) tagged mCherry (**Fig. 1A**) and transduced three human melanoma cell lines – 624-Mel, WM793, and SK-MEL-28. After antibiotic selection, we isolated clonal populations. For each line, clonal growth initially expanded into populations of uniform mCherry expression. Over two weeks in culture, each uniform population yielded bimodal distributions of mCherry expression, such that mCherry low populations begot mCherry high populations and vice versa (**Fig. 1B & Supplemental Fig. S1**). To confirm the fidelity of the reporter we FACS-isolated mCherry low and high cells from each population. The mCherry low cells expressed 2-3-fold less BRN2 protein and mRNA as compared to matched mCherry high cells (**Fig. 1C-D**). Single-molecule (sm)FISH confirmed a significant correlation between the expression of the mCherry transcript and endogenous BRN2 transcript (**Fig. 1E**). Using smFISH, we compared the distributions of BRN2 transcript counts from clonal 624-Mel reporter lines to the parental line (**Fig. 1F**). We detected 3-9 copies of BRN2 mRNA per mCherry low cell, whereas the mCherry high cells presented a broader distribution of 12-60 copies per cell. The distribution of BRN2 transcript within the parental line encompassed the merged distributions of the clones, confirming the clonal populations were representative of the parental BRN2 expression distribution. We performed immunofluorescence for BRN2 in the parental populations and observed similar bimodal expressions (**Fig. 1G**). We generated clonal populations of the parental 624-Mel line via limiting dilution and monitored the BRN2 protein expression over time in culture (**Fig. 1H**). BRN2 expression was uniform in short-term culture and became more heterogenous over time, consistent with the gradual redistribution of mCherry expression we observed in the clones. We conclude that melanoma cells exhibit a heterogenous BRN2 expression pattern that is reestablished upon isolation and clonal outgrowth.

**Figure 1:**
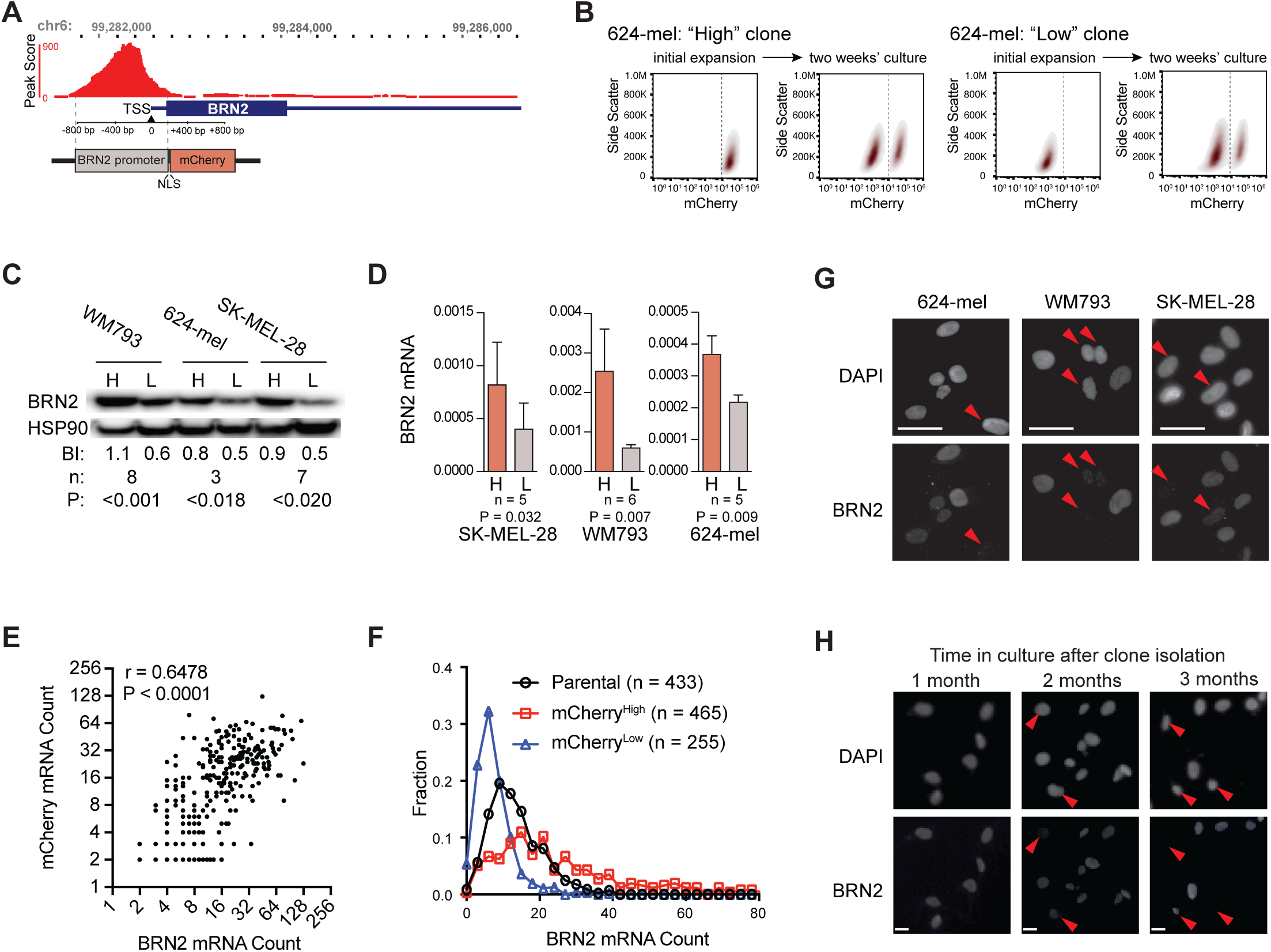
Gradual establishment of BRN2 expression equilibrium in human melanoma cells. A) ATACseq peak scores from the BRN2 locus (top) and schematic of reporter construct (bottom). TSS, transcriptional start site. NLS, nuclear localization signal. B) Flow analysis of clonally expanded then serially cultured 624-mel cells with the BRN2 reporter. See Supplemental Figure S1 for more examples. C) Western blot of BRN2 and HSP90 loading control in reporter high (H) and reporter low (L) cells from indicated cell lines. BI, average blot intensity. n, number of independent lysates. P, two-tailed t test P value. D) Average and SD of ΔCt between probes for BRN2 and RPL37A transcript RT-qPCR probe sets. n, number of independent RNA preparations, P, two-tailed t test P value. E) Single molecule fluorescent in situ hybridization (smFISH) BRN2 and mCherry transcript counts from individual 624-mel cells. r, Pearson’s correlation coefficient. P, two-tailed P value. F) Distribution of BRN2 smFISH transcript counts in 624-mel cells as fraction of total population. n, number of individual cells. G) Images of BRN2 IF staining of indicated cell lines. Red arrowheads indicate cells with poor BRN2 expression. H) Images of BRN2 IF in clonal expansion of 624-mel cells at 1 month, 2 months and 3 months. Scale bars = 20μm.

While culturing, we noticed the cells expressing less mCherry (referred to henceforth as Low^BRN2^) were smaller and more uniform in appearance as compared to the High^BRN2^ cells (**Fig. 2A**). We analyzed isolated Low^BRN2^ and High^BRN2^ cells with digital holographic cytometric feature classification - a method that compares aggregate morphological features of cells^55–57^. In all lines, Low^BRN2^ cells were morphologically distinct from High^BRN2^ cells (P<0.0001) (**Fig. 2B**). We explored phenotypic differences between the lines using live imaging. Low^BRN2^ cells divided more rapidly than High^BRN2^ cells (19.5 hours versus 25.5 hours, respectively, P-value = 0.0028) (**Fig. 2C**). Neither line presented greater random motility (**Fig. 2D**). To gain greater insight into transcriptional changes between the High^BRN2^ and Low^BRN2^ cells, we FACS-purified each population and performed bulk RNAseq. 140 genes were differentially expressed (DE) with Log2 fold change ≥ 1.5 or ≤ -1.5 and adjusted P-value < 0.0005 (**Supplemental Table S1**). Gene set enrichment analysis (GSEA) comparing DE genes to the Hallmark pathways^58^ and to gene sets previously reported in association with melanoma phenotype switching or melanocyte development^13,59^ revealed significant enrichments for signatures of the epithelial to mesenchymal transition, hypoxia, inflammatory response, p53 signaling, undifferentiated melanoma cells, neural crest melanoma cells, and human melanocyte stem cells (MSC) in the Low^BRN2^ population – all pathways previously associated with stem-like tumorigenic cells (**Fig. 2E**). We conclude that clonal human melanoma lines reestablish phenotypic equilibrium in culture between a more stem-like population (Low^BRN2^) and a less stem-like population (High^BRN2^).

**Figure 2:**
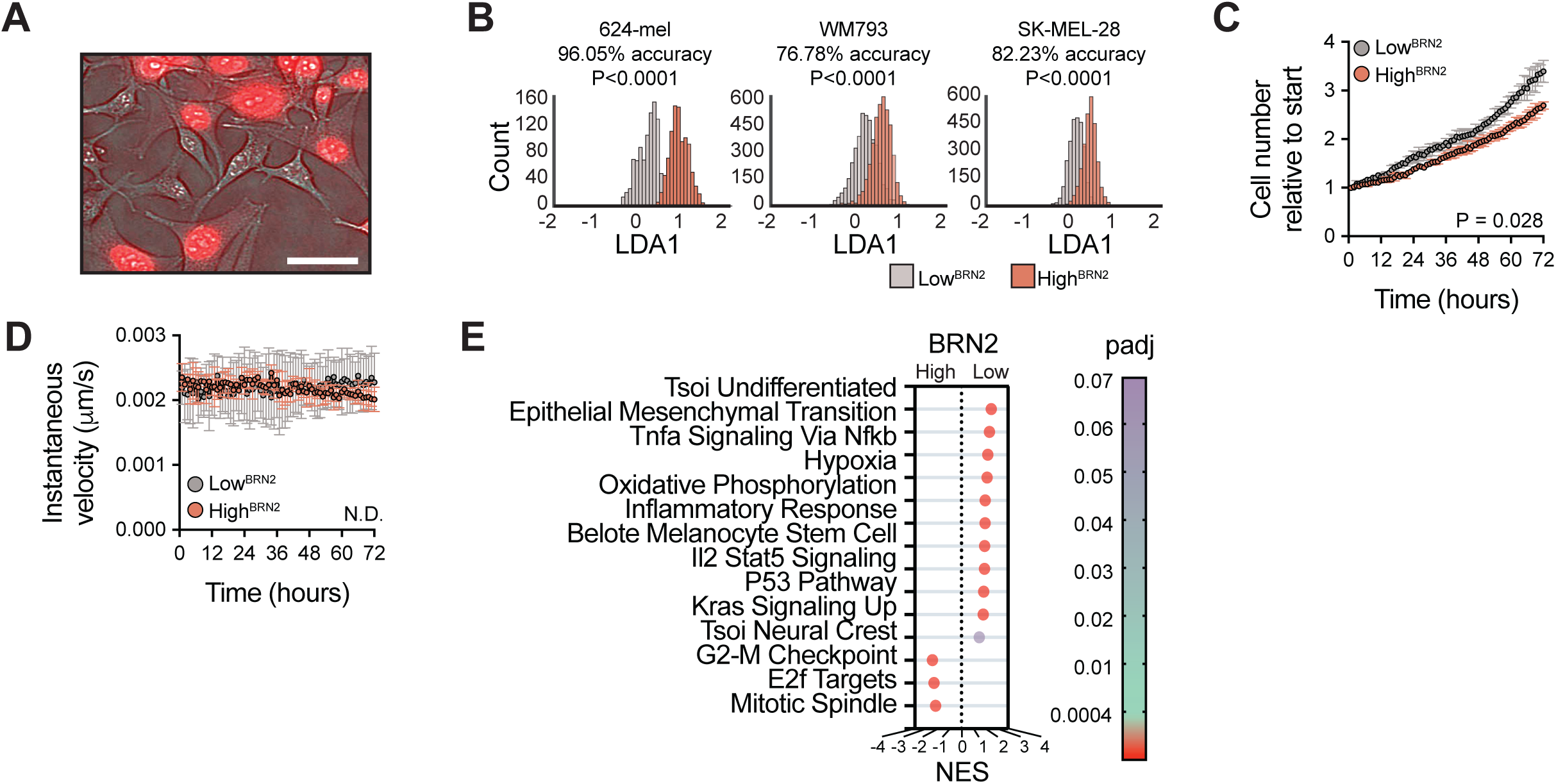
Comparison of phenotype, morphology and gene expression between BRN2 high and low cells. A) Image of clonal 624-mel population expression mCherry reporter. Scale bar = 50μm. B) Classification of indicated cells into mCherry high (High^BRN2^) and mCherry low (Low^BRN2^) populations using a linear discriminant analysis of morphologic features. Population distribution, classification accuracy and p value from two-tailed t test (P) are shown. C) Cell number over time, relative to start of High^BRN2^ and Low^BRN2^ cells. Population mean, standard deviation and p value from two-tailed t test comparing final time point of three independent experiments are shown. D) Instantaneous cell velocity over time from experiments depicted in C). E) Gene set enrichment analysis (GSEA) comparing DE genes between High^BRN2^ and Low^BRN2^ cells to published signatures of melanocyte signaling and differentiation from Belote et al., 2021 and Tsoi et al., 2018 and to Molecular Signature Database Hallmark gene sets. Normalized enrichment score (NES); adjusted p-value (padj).

### Phenotypes are inherited and equilibrium is established via bidirectional interconversion

Evidence for melanoma phenotype switching in prior studies has included the classification of single cells or populations of cells into subgroups based upon both gene expression and cell behavior as well as the emergence of these subgroups in selective conditions, during tumor progression, or upon molecular or genetic manipulation^8,26,29,33^. However, the live monitoring of a single cell undergoing the conversion from one phenotype to another has not been documented in human melanoma cells to our knowledge. As such, the kinetics of phenotype switching in homeostatic conditions, including how switching is related to cell growth and division, remain largely uncharacterized.

We used ptychography^60^ – a form of quantitative phase imaging – coupled with fluorescent microscopy to monitor heterogenous 624-mel cultures expressing the mCherry reporter. Cells were monitored for 48 hours at 30-minute intervals. We tracked the lineages of cells identified at timepoint zero and monitored mCherry expression level and cell morphology (**Fig. 3A-C**).

**Figure 3:**
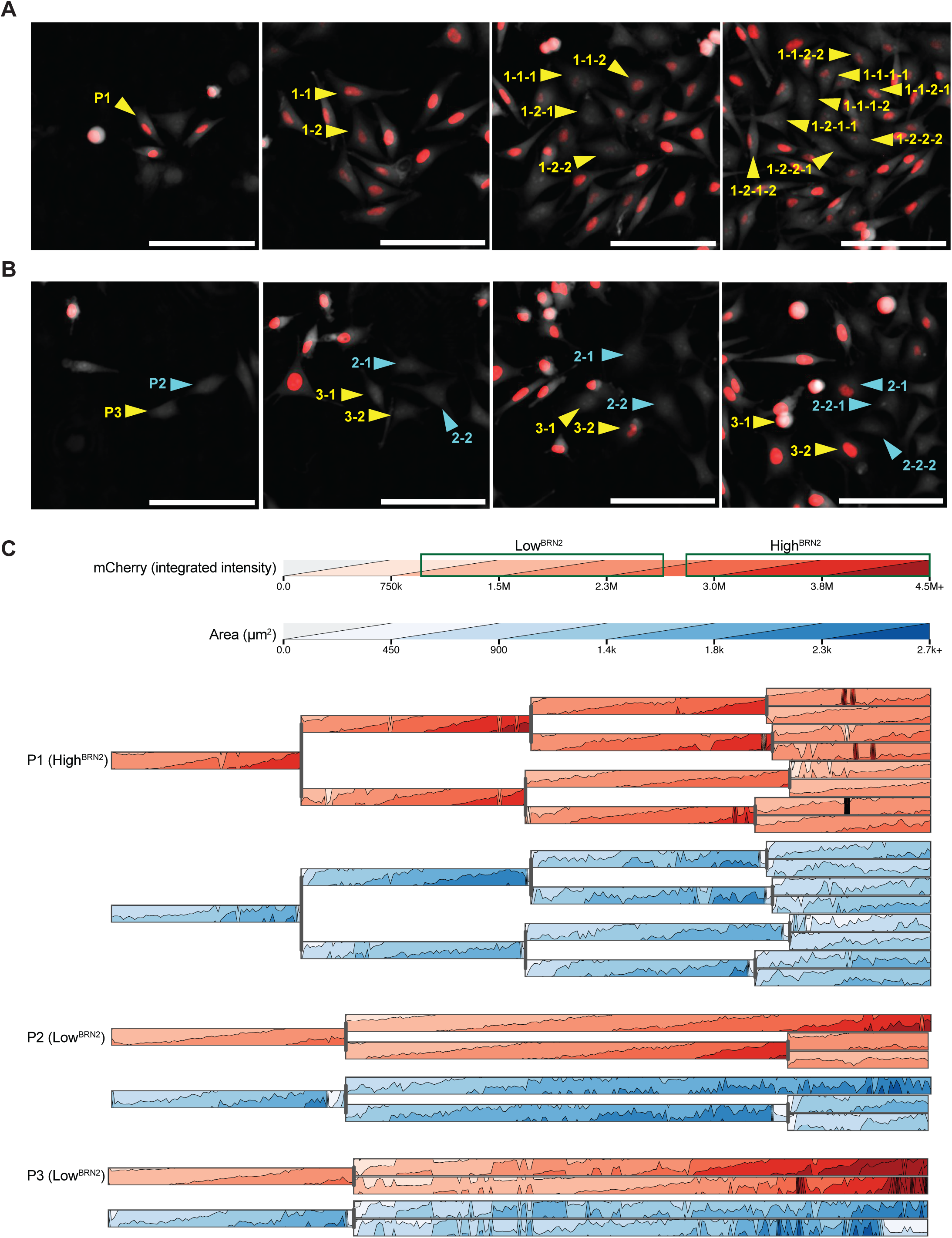
Phenotypes are both inherited and undergo spontaneous bidirectional interconversion. A-B) Stills from representative Supplemental Videos 1 and 2, respectively, taken every 30 minutes for 2 days. Parental cells (P1, P2, P2) are highlighted in each (left column) and tracked through one or more divisions (yellow and blue arrow heads). Scale bar = 200μm. C) Integrated mCherry intensity and cell area over time of lineages highlighted in (A), plotted as lineage tracking horizon charts. Green rectangles indicate range of mCherry expression during cell growth and division for High^BRN2^ and Low^BRN2^ cells. A visual key to horizon chart interpretation is in Supplemental Figure S2.

Overall, mCherry expression was stable in most cells over multiple divisions (**Supplemental Videos V1-V2**). Expected fluctuations of mCherry that corresponded to the accumulation and subsequent halving of cell mass associated with growth and division were observed with each cell cycle, but these changes in mCherry expression were minor in comparison to the difference in mCherry expression between the two states (**Fig. 3C**). During most divisions, the mCherry expression level of the parental cell was inherited by each daughter cell, resulting in clusters of either High^BRN2^ or Low^BRN2^ cells. Interconversions in both directions were also captured. For example, we observed High^BRN2^ parental cells giving rise to both High^BRN2^ and Low^BRN2^ daughter cells (**Fig. 3A**, cells 1-1, 1-1-1, and 1-1-2) and the subsequent reverse transition of Low^BRN2^ progeny back to a High^BRN2^ phenotype (**Fig. 3A**, cells 1-1-1-1 and 1-2-1-2). In many cases, the transitions occurred as apparent asymmetric divisions, although we also observed the gradual increase of mCherry expression in Low^BRN2^ cells outside the context of division (**Fig. 3B**, cell 3-2). The changes in mCherry expression were coupled with the expected changes in cell morphology, with mCherry high cells adopting a larger and flatter morphology (**Fig. 3C**, area), further demonstrating the fidelity of the reporter in marking distinct cell phenotypes. These observations confirm that phenotype switching occurs in human melanoma cells in homeostatic conditions. The two phenotypes can be inherited and typically remain stable through multiple cell divisions, though sporadic bidirectional phenotype switches do occur.

### The Low^BRN2^ phenotype is tumorigenic

We next investigated whether phenotypes associated with either Low^BRN2^ or High^BRN2^ cells might impact melanoma growth or metastatic behavior. One phenotype associated with aggressive disease is the ability to initiate tumorigenesis or metastasis upon engraftment into mice^61^. The enrichment for neural crest, MSC, hypoxia and other similar programs in the Low^BRN2^ cells (**Fig. 2E**) is consistent with a stem-like phenotype^30,32^. We, therefore, tested tumorigenicity using a graft efficiency assay. Reporter 624-mel clones were transduced with a constitutively active luciferase construct. Two million FACS-enriched Low^BRN2^ or High^BRN2^ cells were isolated via FACS and implanted subcutaneously into NOD *scid* gamma mice. Tumor growth was monitored over five weeks (**Fig. 4A**). Mice implanted with Low^BRN2^ cells formed tumors in 100% (7/7) of mice compared to 14.3% (1/7, P = 0.0018) of mice implanted with High^BRN2^ cells (**Fig. 4B-C**).

**Figure 4:**
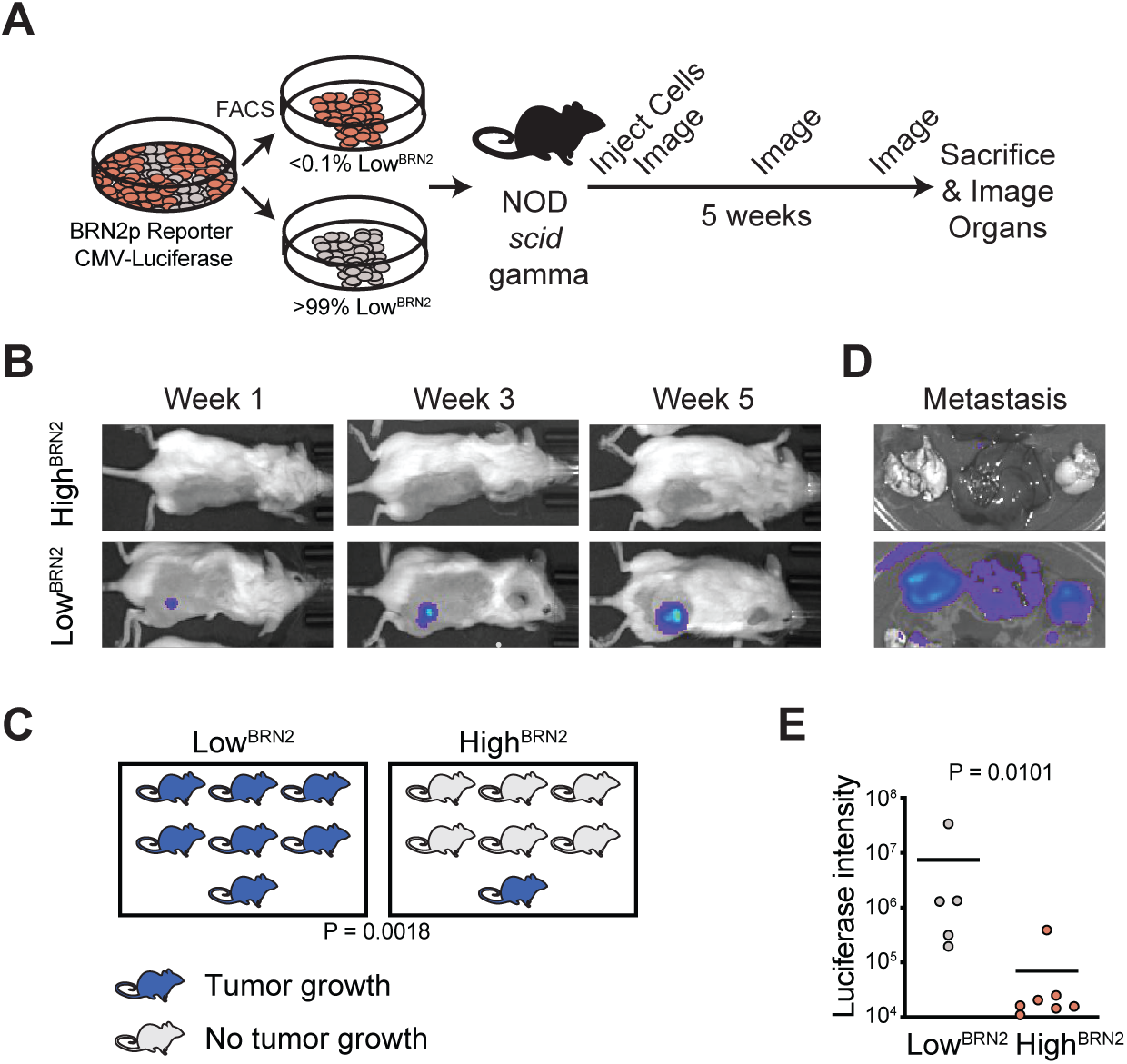
The BRN2 low state is tumorigenic. A) Schematic of graft efficiency assay. Clonal 624-mel cultures were either depleted (red) or enriched (grey) for Low^BRN2^ cells and implanted into NOD scid gamma mice. Luciferase imaging was conducted 1, 3, and 5 weeks after implantation followed by sacrifice and organ imaging. B) Representative images of luminescence at indicated time points. C) Percent of mice with successful grafts after 5 weeks. P, two-tailed Fisher’s exact test P value with 0.98 power. D) Representative images of organ luminescence. E) Average (line) and individual summations of luciferase intensity in brain, liver, and lungs. P, two-tailed t test P value.

After five weeks, surviving mice were sacrificed, and the brain, lungs, and liver were imaged (**Fig. 4D**). All mice engrafted with Low^BRN2^ cells presented distal metastatic dissemination based on luciferase expression, as compared to a single mouse in the High^BRN2^ cohort (P = 0.0101) (**Fig. 4E**). Thus, Low^BRN2^ cells were more competent for initiating tumor growth and metastasis upon engraftment.

### Tolerance to targeted BRAF^V600E^ inhibition excludes the Low^BRN2^ phenotype

In addition to primary growth and metastasis, another phenotype associated with aggressive disease is resistance to therapeutic intervention. Each of the three lines selected for this study harbors the *BRAF^V600E^* oncogenic driver, identified in approximately half of all melanomas^62^ (confirmed by digital droplet RT-qPCR, data not shown). Small molecules that selectively inhibit the mutated BRAF^V600E^ kinase, including vemurafenib, dabrafenib and encorafenib, are the predominant targeted kinase therapies used to treat *BRAF^V600E^* positive melanomas, but frequently result in widespread treatment resistance^63^. We FACS-isolated Low^BRN2^ and High^BRN2^ 624-mel cells and conducted vemurafenib response studies. We first used ptychography coupled with fluorescence microscopy to track the accumulation of cell mass over time – a sensitive readout of changes in cell growth in response to drug^64^. At high doses (1-10 μM) exposure to vemurafenib inhibited cell mass accumulation in both populations (**Fig. 5A**). Interestingly, at low concentrations (0.1 μM), we observed a paradoxical increase in mass accumulation exclusively in High^BRN2^ cells (P < 0.0001). We further noticed that in both populations, despite the decrease in mass accumulation at high doses, exposure to vemurafenib induced an increase in mCherry expression (**Fig. 5B-C**). At the population level, FACS-isolated Low^BRN2^ populations were more sensitive to the compound, exhibiting an IC50 of 0.26μM (95% CI: 0.15-0.45), consistent with the mass measurements (**Fig. 5D**). In contrast, the High^BRN2^ populations presented a more complex response to vemurafenib. Specifically, at low doses (0.001-0.010 μM) vemurafenib again induced a minor, but non-significant, increase in cell number over 36 hours of treatment (**Fig. 5D**). At higher doses the High^BRN2^ populations exhibited an IC50 almost 10-fold higher than that of Low^BRN2^ cells (2.15μM, 95% CI: 0.73-6.37, P < 0.0005, unpaired t test). Flow analysis confirmed that exposure to vemurafenib induced a phenotype transition, such that all surviving cells uniformly expressed increased mCherry, thus depleting the population of Low^BRN2^ cells (**Fig. 5E**). Coupled with the observed necessity of the Low^BRN2^ state for graft efficiency (**Fig. 4**), our experiments demonstrate that BRAF^V600E^-driven melanoma cells interconvert between two mutually exclusive and inheritable states – each of which promotes a distinct behavior associated with aggressive disease.

**Figure 5:**
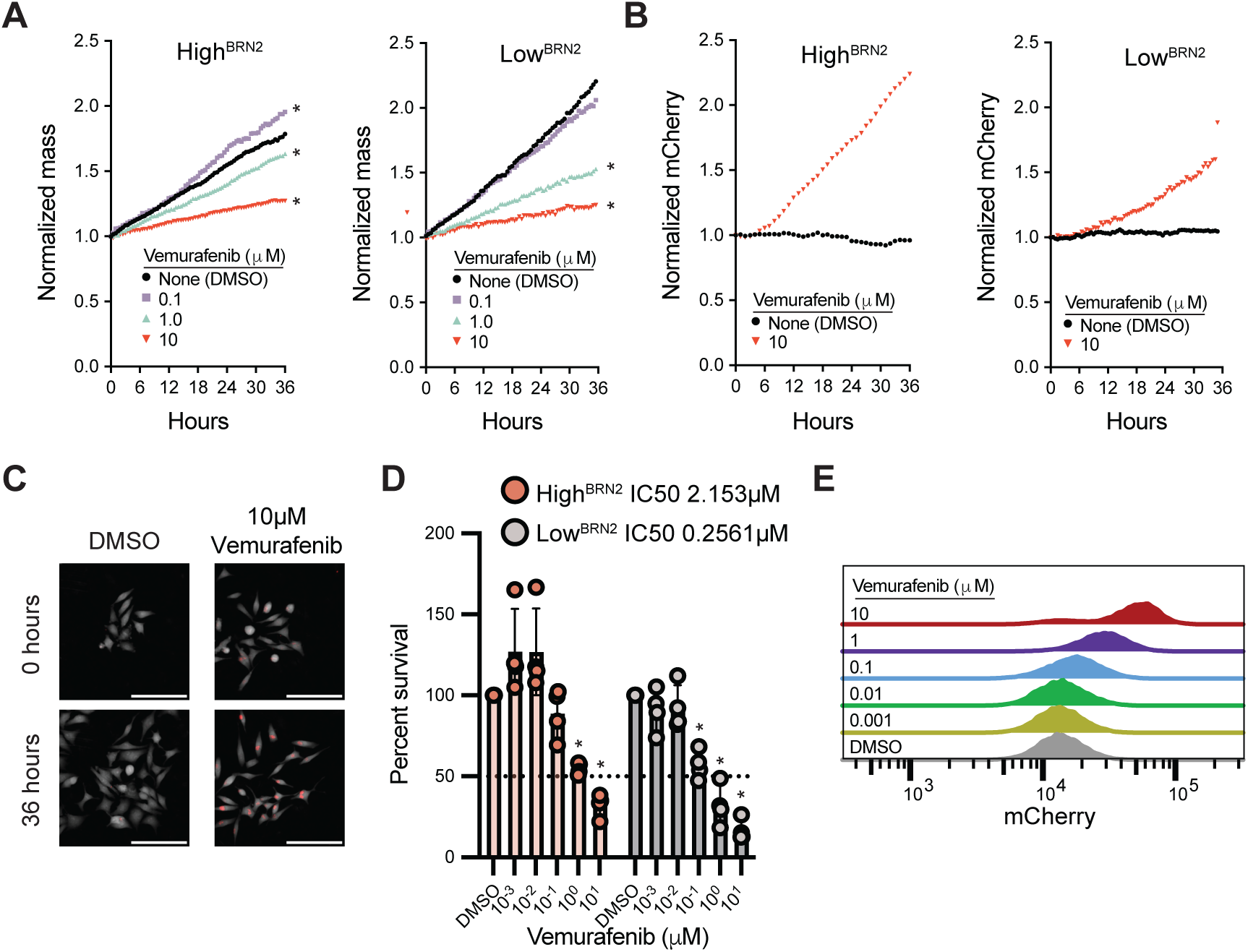
Tolerance to targeted BRAF^V600E^ inhibition excludes the Low^BRN2^ phenotype. A) Normalized mean single cell mass over time of FACS-enriched High^BRN2^ (left) and Low^BRN2^ (right) 624-mel cells exposed to indicated concentrations of vemurafenib. Asterisk indicates slopes significantly different from DMSO condition (P < 0.0005, simple linear regression). B) Normalized mean integrated mCherry intensity over time of FACS-enriched High^BRN2^ (left) and Low^BRN2^ (right) 624-mel cells exposed to indicated concentrations of vemurafenib. C) Representative image of Low^BRN2^ cells from B. Scale bar = 200μmt. D) Percent surviving cells after 48 hours of exposure to indicated concentrations of vemurafenib relative to DMSO condition. Asterisk indicates means significantly different from DMSO condition (P < 0.0005, unpaired T test). E) Flow analysis of mCherry expression from 624-mel reporter cells after 48 hours treatment with the indicated concentrations of vemurafenib.

### BRN2-associated drug-tolerant and tumorigenic phenotypes are present in human melanoma

We performed single-cell RNA sequencing (scRNAseq) on FACS-isolated Low^BRN2^ and High^BRN2^ 624-mel cells to explore the similarity and differences of individual cells within each population. Uniform Manifold Approximation and Projection (UMAP) revealed that while Low^BRN2^ and High^BRN2^ generally clustered together, most cells assembled as a single large population (**Fig. 6A, Supplemental Figure S3A**). Seurat clustering subdivided the population into nineteen clusters, each with varying fractions of Low^BRN2^ and High^BRN2^ cells (**Fig. 6B, Supplemental Figure S3B**). After examining previously published gene expression programs regulated by BRN2 and three other lineage-specific factors, MITF, AXL and SOX10, known for their roles in phenotype switching and targeted therapy response^17,34,35^, we classified Seurat clusters into three groups (**Fig. 6C-D, Supplemental Figure S3C-D**). Cluster 5, composed of almost exclusively High^BRN2^ cells (**Supplemental Figure S3B**), had the highest expression of BRN2-regulated genes (“BRN2 genes”)^26^ and AXL-regulated genes (“AXL genes”)^16^ (**Fig. 6C, E, F**) and so we refer to it as the BRN2 state (**Fig. 6D**, green). Notably, the BRN2 state exhibited poor expression of both SOX10-regulated genes (“SOX10 genes”)^45^ and MITF-regulated genes (“MITF genes”)^16^ (**Fig. 6C, E-H, Supplemental Figure S3C-D**). Conversely, clusters 0 and 6 were comprised mostly of Low^BRN2^ cells (**Supplemental Figure S3B)**, expressed less BRN2, AXL and MITF genes, but retained high expression of SOX10 genes (**Fig. 6C,E-H, Supplemental Figure S3C-D**), and so we refer to these as the SOX10 state (**Fig. 6D**, red). Finally, all other clusters, regardless of reporter status, expressed a similar pattern of uniformly highly expressed MITF genes, mediumly expressed SOX10 genes, and poorly expressed BRN2 and AXL genes (**Fig. 6C, E-H, Supplemental Figure S3C-D**), and instead differed from each other by cell cycle genes (**Supplemental Figure S3E-F**). We collectively refer to these as the MITF state (**Fig. 6D**, blue). We next examined the fractional composition of each group in the bulk population of Low^BRN2^ and High^BRN2^ cells. As expected, BRN2 state cells were near-exclusive to the High^BRN2^ population and SOX10 state cells were near-exclusive to the Low^BRN2^ (**Fig. 6I**). Consistent with these data, SOX10 protein expression was reduced in the High^BRN2^ populations of all reporter lines (**Fig. 6J**). Our identification of a targeted therapy resistant SOX10 deficient population of melanoma cells, which exhibit relatively slower proliferation (**Fig. 2C**), is also consistent with previous reports^34,65^. Further validating the BRN2 state as the targeted therapy resistant cell state previously reported, this state was enriched for TGF-beta and EGFR gene expression programs (**Fig. 6K**) and bulk High^BRN2^ cells expressed higher protein levels of EGFR relative to Low^BRN2^ cells (**Fig. 6L**).

**Figure 6:**
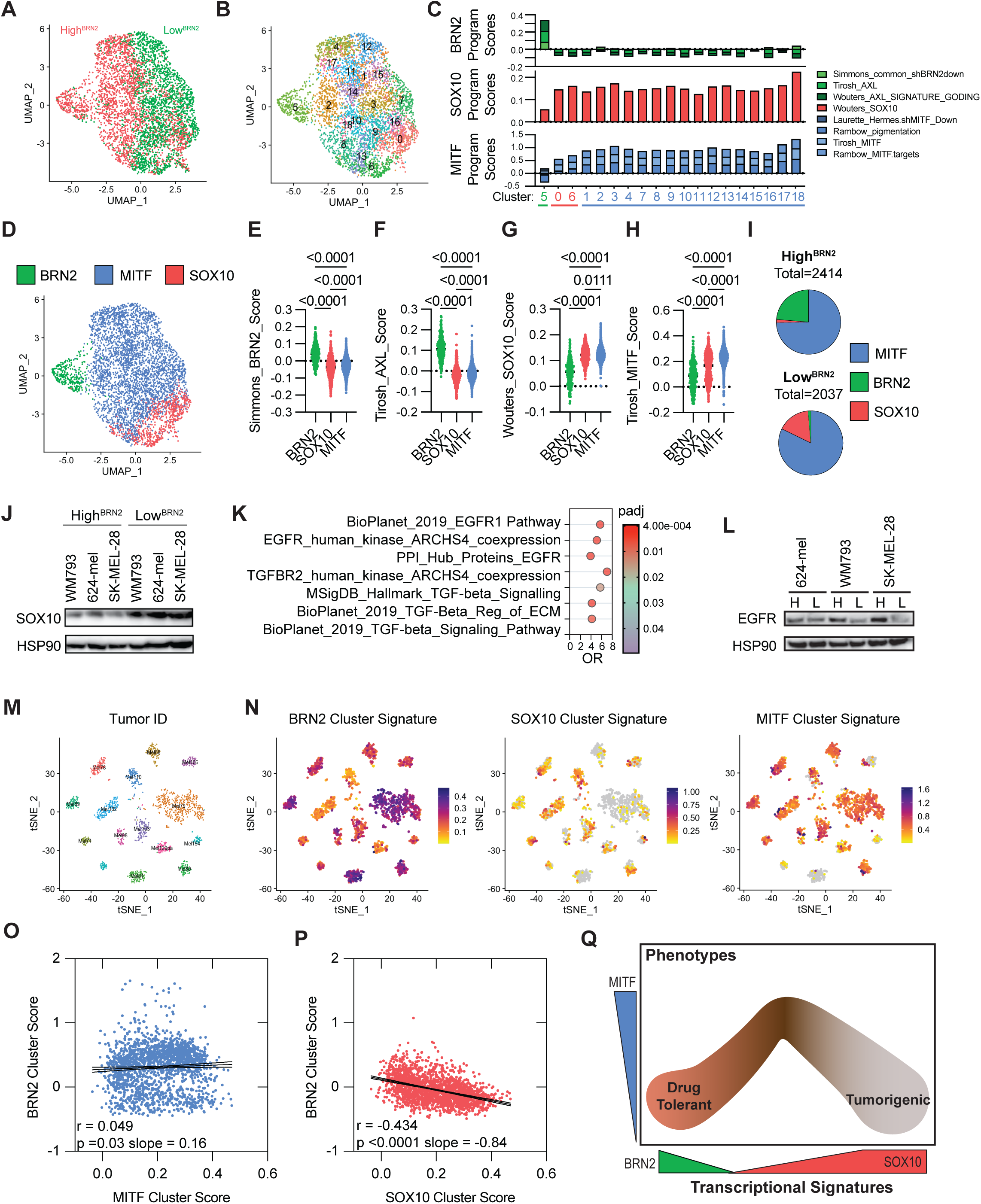
Distinct and mutually exclusive MITF low transcriptional states in human melanomas. A) Uniform manifold approximation and projection (UMAP) visualization of the 4,451 cells that passed quality control (Supplemental Fig. S3A), colored according to FACS enrichment (High^BRN2^ or Low^BRN2^). B) UMAP colored according to Seurat clustering (see also Supplemental Fig. S3B). C) ModuleScores for previously published BRN2 and AXL regulated genes (green); SOX10 regulated genes (red) and MITF regulated genes (blue). D) UMAP colored according to BRN2, SOX10, and MITF clusters. E-H) Distribution of gene signature scores within cells of the BRN2, NCSC, and MITF clusters (see also Supplemental Fig. S3C-D for UMAP representations of signature scores). P, ordinary one-way ANOVA. I) Pie charts indicating the percent of High^BRN2^ and Low^BRN2^ cells from the BRN2, SOX10, and MITF clusters. J) Western blot analysis of SOX10 expression in FACS enriched mCherry high and mCherry low reporter cell lines. K) Gene set enrichment analysis (GSEA) comparing DE genes between High^BRN2^ and Low^BRN2^ cells to signatures of known signaling pathways. L) Western blot analysis of EGFR expression in FACS enriched mCherry high and mCherry low reporter cell lines. M) tSNE plots of ex vivo human melanoma scRNAseq data from Jerby-Arnon, 2018. Tumor IDs identify unique patients. N) Scores for gene signatures unique to BRN2, SOX10, and MITF clusters are plotted as colorimetric onto tSNE plot from (M). O) MITF and N) SOX10 cluster scores versus BRN2 cluster scores for every cell in (M). Best-fit line, 95% CI from simple linear regression; r and p, Pearson correlation. N) Schematic summary of the relationship between observed transcriptional signatures (green, red, blue) and observed phenotypes (drug-tolerant and tumorigenic).

We next aimed to ascertain if the triad of states delineated via scRNAseq and this reporter system were reflective of transcriptional conditions present within human tumors. For each cluster, we identified a distinct signature defined by a unique combination of differentially expressed genes (**Supplemental Table S2**). These distinct signatures were subsequently evaluated in a scRNAseq dataset encompassing fourteen *ex vivo* human melanoma tumors (**Fig. 6M**) generated by Tirosh et al^16^ and Jerby-Arnon et al^66^. Consistent with previous reports, most tumors were comprised of cells heterogeneously expressing the MITF signature, the BRN2 signature, and/or the SOX10 signature (**Fig. 6N**)^14,16,34^. Interestingly, the BRN2 state signature was expressed in a largely bimodal manner – with distinct high and low expression states (**Fig. 6O**) not unlike the BRN2 promoter reporter expression pattern (**Fig. 1B**). While there was no obvious relationship between BRN2 and MITF state signature expression (**Fig. 6O**), the BRN2 and SOX10 states presented a significant anticorrelated expression pattern (**Fig. 6P**, r = -0.434, p <0.0001). These findings reinforce that the mutually exclusive transcriptional programs delineated in our reporter system aptly reflect the conditions present within human melanoma tumors.

## Discussion

In this study, we used label-free live cell imaging coupled with a fluorescent reporter of BRN2 promoter activity to monitor the kinetics of phenotype switching in human melanoma cells in homeostatic conditions. Our work builds upon prior studies that either indirectly monitored phenotype switching by taking molecular snapshots at different time points or induced the switch through chemical, environmental, or genetic perturbation^21^. Through tracking individual live cells, we demonstrate phenotype heterogeneity within clonal populations that is re-established in standard culture conditions, as has been previously demonstrated for breast cancer and glioblastoma^67,68^. Switching occurs on an intermediate time scale, such that phenotypes are inherited through multiple generations while also undergoing spontaneous interconversion^69^. Transitions occurred both coupled with asymmetric division or independent of cell division. Since switching was bidirectional, our observations reinforce the dynamic stemness model in melanoma^1^ where stem-like and non-tumorigenic cells reach a phenotypic equilibrium, potentially leading to unlimited tumorigenic potential if the stem-like state cannot be controlled. Further investigations into the underlying mechanisms that facilitate dynamic stemness and ways to prevent this transition are needed.

Pioneering work in this domain predicted that melanoma cells switched between two phenotypes – a MITF high and BRN2 low proliferative state and a MITF low and BRN2 high invasive state^15^. Our approach yielded observations consistent with recent studies that demonstrate a more complex phenotypic landscape within populations of melanoma cells^14,22,26,31,32,39,45^. Notably, we observed neither an obligatory relationship between BRN2 expression and migration nor evidence for a MITF-BRN2 axis. Instead, we observed two distinct MITF low transcriptional states characterized by the expression of either a BRN2- or a SOX10-regulated signature and associated with mutually exclusive phenotypes of drug tolerance and tumorigenicity (**Fig. 6Q**). The observed coexistence of these three states, within not only the isogenic and clonal homeostatic culture but also within human tumors, provides a plausible reconciliation for prior inconsistent findings in the field concerning the relationship between MITF and BRN2 programs. For instance, cultures classified as "MITF low" may actually constitute a blend of BRN2 and SOX10 states, with proportions potentially varying or even oscillating. In a similar vein, our data suggest that "SOX10 low" cultures are likely to exhibit a uniformity of low MITF levels, whereas "SOX10 high" cultures could encompass cells with a blend of high and low MITF. Further, this cell state heterogeneity in cultures and tumors would inevitably impact observed phenotypes, especially when only population means are considered. The two low MITF states, which reciprocally express BRN2 and SOX10, are likely akin to the BRN2 and NCSC states identified in earlier studies^1,14,31,34,70^. Consistently, SOX10 low melanoma cells (identified as High^BRN2^ in this study) have been associated with increased tolerance to targeted therapy, whereas SOX10 high cells (Low^BRN2^ in this study) were depleted in drug-resistant tumors^14,34^. The mutual exclusivity of these two states, yet their individual contribution to different stages of melanoma progression (i.e., tumor initiation versus therapeutic resistance), offers a rational explanation for the paradoxical identification of BRN2 as both an oncogene and a tumor suppressor^8,23,33^.

It is noteworthy that our single-cell phenotyping analysis suggested more uniformly discrete cell populations than what the scRNAseq analysis implied. This finding points towards the possibility that variations in the melanoma transcriptome, observed as distinct clusters in scRNAseq, do not invariably cause observed alterations in the phenome. This lack of supervenience of the transcriptome on the phenome aligns with both the broadly recognized general mismatch between transcript and protein expression^71,72^, as well as the specific, complex layers of post-transcriptional regulation of MITF and BRN2^25^. It is also in line with the overarching phenomenon of biological robustness within cells^73,74^. This insight underscores the potential shortcomings of relying solely on scRNAseq to infer cell phenotypes. As strides are made toward characterizing distinct melanoma phenomes, the potential disconnect between the transcriptome and phenotype should be considered.

Prior studies have demonstrated that BRAF inhibitor-resistant cells also exhibit increased invasive or stem-like properties – linking the two deleterious phenotypes^70,75^. We were therefore surprised to identify distinct aggressive phenotypes associated with each mutually exclusive state. Our data suggest that once a melanoma is established, the induction of either state could progress the disease in discrete ways – increasing metastatic dissemination or decreasing therapeutic sensitivity. One limitation of our study is its reliance on melanoma cell lines for assaying phenotypes. Although we present evidence of both states existing within human tumors, we are unable to conclude that either state is associated with the same behaviors in situ as we observed in vitro and in vivo. We hypothesize that the burden of BRN2 state cells in patients’ tumors could be predictive of the rate at which targeted therapy resistance occurs. Conversely, the percentage of SOX10 state cells could be prognostic of the likelihood of widespread dissemination or remission. The assembly of appropriate cohorts for assessing these hypotheses should be the subject of future investigation.

Overall, this research provides useful insights into the dynamics of phenotype switching in melanoma cells. Our findings highlight the importance of understanding the mechanisms underlying phenotypic switching in melanoma and the relationship between therapy-tolerant and stem-like tumorigenic properties. In particular, we emphasize the implications of our observations for strategies seeking to enhance the efficacy of melanoma therapies by targeting the mechanisms that control phenotype switching. As the induction of either phenotype could be detrimental to a patient, future studies should weigh the potential benefit of sensitizing cells to therapeutic regimens against the potential harm of increasing the risk of metastatic outgrowth.

## Materials and Methods

### Cell Culture Maintenance and Manipulation

Melanoma cell lines 624-Mel (CVCL_8054), SK-MEL-28 (CVCL_8054) and WM793 (CVCL_8787) were received as gifts from Dr. Boris Bastian and Dr. Meenhard Herlyn and were STR verified (**Supplemental Table S3**) and regularly monitored with the Universal Mycoplasma Detection Kit (ATCC 30-1012K). Cultures were maintained in RPMI1640 media (Thermo Fisher, 11-875-093) supplemented with 10% FBS (Corning, 35-010-CV), 1x penicillin-streptomycin, (Thermo Fisher, 15140122) and 2mM L-glutamine (UCSF core facility, CCFGB002) at 37 °C, 5% CO_2_. All cells were disassociated with 0.05% Trypsin EDTA (Mediatech, MT 25-052-C1) followed by neutralization with equimolar soybean trypsin inhibitor (Thermo Scientific, 17075029). The BRN2 reporter (pROM-POU3F2p-mCherry-Neo, Addgene 153321) was engineered by first inserting NLS-mCherry (Gift from Ron Vale, Addgene 67932) and a pGK-driven neomycin resistance cassette into a 3^rd^ generation vector backbone, pSicoR-Ef1a-mCh-Puro, (Gift from Bruce Conklin, Addgene 31845)^76,77^. The BRN2 promoter was amplified off of human DNA (**Supplementary Table S4**). Lenti-viral particles were generated as previously described and transduced at MOI < 0.3 in the presence of 10 μg/mL polybrene (Sigma TR-1003)^23^. Three days later, pROM-POU3F2p-mCherry-Neo transduced cells were selected with neomycin (Sigma, A1720) for 3 weeks at the minimal concentration required for complete toxicity of parental cells (cell line dependent, concentrations ranged from 100 μg/ml to 1000 μg/ml). Single cells from each selected culture were sorted using the SONY SH800 FACS machine and expanded. Neomycin was periodically added to the media to ensure retention. For isolation experiments, respective populations were FACS isolated twice in tandem followed by a third analyses to ensure >99% purity. Flow analysis were conducted with a BD Fortessa.

### ATAC-Seq

The ATAC-seq assay followed the published Omni-ATAC protocol^78^. Briefly, nuclei were isolated from 50000 cells, which were then treated with 2.5ul of Tn5 transposase (Illumina 20034197) for DNA tagmentation. DNA was extracted and PCR amplified (5 cycles) using barcoded primers that were included in the Omni-ATAC protocol^78^. The ATAC libraries were sequenced by NovaSeq (50bp paired end). ATAC-sequenced reads were quality checked by FastQC (v0.11.9), and then aligned to the human reference genome hg19 (GRCh37) using Bowtie2 aligner (bowtie2-align-s version 2.4.2). Utilizing samtools (version 1.12), all mitochondrial, unaligned, and low-quality reads (Q<20) and duplicated reads were filtered out, and were then normalized for equal coverage in each group of cells. Removing adapters by NGmerge (version 0.3), ATAC-enriched regions/peaks were identified by MACS (3.0.0a6) with Q value cut off of 0.01 and with other default parameters. The identified peaks were visualized through Integrated Genome Browser (IGB), and also were annotated by HOMER (v4.11.1-the annotatePeaks.pl program). Finally, the differentially enriched peaks between two groups of cells were evaluated via DiffBind (version 3.4.3), using DESeq2 method.

### Protein preparation and Western blotting

At least 200,000 cells were lysed in RIPA buffer (Thermo 89901) containing a protease inhibitor cocktail (Thermo 87785) and 0.5 M EDTA (Thermo 1861274), each used at 1:100 concentration, incubated on ice for 10 minutes, then centrifuged at maximum speed for 20 minutes at 4°C. The protein concentration of the supernatant was measure using the Pierce BCA protein assay kit (Thermo 23225) and 10 μg of protein was loaded on a 4–12% Nupage bis-tris gel (Thermo NP0323box), run for 2 hours at 130 V in running buffer (Novex NP0002), and dry-transferred onto a nitrocellulose membrane (Biorad 1620115) using a Bio-Rad Trans-blot SD semi dry transfer cell. The membrane was submerged in blocking buffer (Thermo, 37536) for 1 hour before incubation with a primary antibody diluted in blocking buffer (**Supplemental Table S4**) overnight, followed by trice washing in 0.2% Triton-X (Sigma T8787) in PBS before a secondary antibody (Invitrogen G-21040 or Invitrogen G-21234, 1:2000) was added and incubated for 1 hour. After three washes, the membrane was imaged using an Azure Biosystem Imager. The band intensity was measured by densitometry using Image J 1.52a.

### RNA preparation, and quantitative real-time PCR, bulk RNA sequencing

FACS isolated cells were added to 0.5-1 mL of Trizol (Ambion 15596026) for RNA extraction according to manufacturer’s protocol. For qRT-PCR, cDNA from 0.5-1 μg of RNA (NanoDrop, Thermo Scientific) was synthesized using the Sensifast synthesis kit (Bioline Bio-65053, manufacturer’s protocol) then diluted 1:5. Per RT-qPCR reaction, 1 μl of the dilution was mixed with 0.41 μM primers (**Supplemental Table S4**) and 1x Sensifast SYBR non-Rox kit (Bioline Bio98005). The real-time PCR program cycled as follows: one cycle of 95°C for 2 mins followed by 40 cycles of 95°C for 5 seconds and 65°C for 30 seconds. For sequencing, intact poly(A) RNA was purified from 100-500 ng of RNA (RIN 8-10, Agilent Technologies 2200 TapeStation) with oligo(dT) magnetic beads and stranded mRNA sequencing libraries were prepared as described using the Illumina TruSeq Stranded mRNA Library Prep kit (20020595) and TruSeq RNA UD Indexes (20022371). Purified libraries were qualified on an Agilent Technologies 2200 TapeStation using a D1000 ScreenTape assay (cat# 5067-5582 and 5067-5583). The molarity of adapter-modified molecules was defined by quantitative PCR using the Kapa Biosystems Kapa Library Quant Kit (cat#KK4824). Sequencing libraries (1.3 nM) were chemically denatured and applied to an Illumina NovaSeq flow cell using the NovaSeq XP chemistry workflow (20021664). Following transfer of the flowcell to an Illumina NovaSeq instrument, a 2 x 51 cycle paired end sequence run was performed using a NovaSeq S1 reagent Kit (20027465). Gene set enrichment analyses were conducted as previously described^7^.

### Single molecule RNA FISH

smFISH experiments were conducted with in house reagents as previously described^79^. Briefly, probes were developed using the designer tool from Stellaris (LGC Biosearch Technologies) using a masking level of 5, a minimum of 2 base pair spacing between single probes, and a length of 18 nt (**Supplemental Table S4**). Approximately 5×10^5^ disassociated cells were immobilized on a Cell-Tak (Corning, CB-40240) coated 8-well chambered image dish, fixed with 5% formaldehyde (Tousimis 1008A), 1X PBS for 10 minutes, washed with 1X PBS then stored in 70% EtOH at 4 °C for a minimum of one hour. Cells were then washed first with 2X SSC (Thermo Fisher Scientific 15557044) containing 10% v/v deionized Formamide (Thermo Fisher Scientific AM9342), then with hybridization buffer consisting of 10% w/w dextran sulfate (Sigma Aldrich 42867) in 1X SSC with 10% v/v Formamide, and incubated overnight at 37 °C in hybridization buffer containing 25 nM probes. Cells were then incubated for 30 minutes at 37 °C in 2X SSC containing 10% v/v Formamide, followed by 15 minutes at 37 °C in 2X SSC containing 10% v/v Formamide and DAPI (Thermo Fisher Scientific D1306). Finally, cells were washes with 2X SSC and incubated for 2 min at room temperature in 2X SSC, 10% w/w glucose (Sigma Aldrich G7021), 0.01 M Tris pH 8 (Life Technologies AM9855G). To minimize photo bleaching, cells were imaged in a photo-protective buffer containing 2X SSC, 10% w/w glucose, 0.01 M Tris pH 8, 75 μg/mL glucose oxidase (Sigma Aldrich G0543-10K), 520 μg/mL catalase (Sigma Aldrich C3156-50), and 0.5 mg/mL Trolox (Sigma Aldrich 238813). Images were taken with a Nikon Ti-E microscope equipped with a W1 Spinning Disk unit, an Andor iXon Ultra DU888 1k x 1k EMCCD camera and a Plan Apo VC 100x/1.4 oil objective in the UCSF Nikon Imaging Center. Approximately 10 xy locations were randomly selected for each condition, and analyzed using Fiji and in-house programs^79,80^.

### Quantitative Immunofluorescence

Cells were fixed in 4% paraformaldehyde (PFA) in PBS (Biotium 22023) for 15 minutes at room temperature followed by pre-cooled (-20°C) methanol for 10 min at -20°C. Cells where thrice washed in PBS then incubated in blocking buffer (5% Donkey Serum (Jackson Immuno Research Labs 017-000-121, 1% BSA (Sigma A9647), 0.1% Triton X-100 (Sigma T8787) in PBS) for one hour at room temperature. Cells were incubated at 4°C overnight with anti-BRN2 (Cell Signaling 12137S, 1:1000), thrice washed in PBS and incubated at room temperature for 1 hour with goat anti-rabbit Alexa Fluor Plus 488 (Invitrogen, A32731, 1:10000). Cells were washed, incubated with DAPI (Thermo Fisher D1306, 1:1000) for 5 min at room temperature, washed twice more and imaged using the INCell Analyzer 2000 high throughput high content imager (GE).

### Quantitative Phase Imaging to Assess Proliferation, Motility, Morphology and Reporter Fluorescence

100,000 cells were seeded per well of a standard tissue-culture treated 6-well polystyrene plate (Sarstedt 83.3920.005) for digital holographic cytometry (DHC) using the M4 Holomonitor (Phase Holographic Imaging, Sweden). Cells were monitored for 72 hours. Morphology was analyzed using the HStudio Software and linear discriminant analysis as previously described^55–57^. For Fourier pthychography coupled to fluorescent microscopy, 1,200 cells were seeded per well of a standard-culture treated 96-well plate then imaged with the LiveCyte (Phasefocus, United Kingdom) every 30 minutes for 48-72 hours. Proliferation and motility were analyzed using the Analysis dashboards provided by the manufacturer. Cell mass was calculated by segmenting cell objects from background using Sobel edge detection thresholding, then summing the phase shift relative to background over all object (cell) pixels, and assuming a specific refractive increment of 1.8×10^-4^ m^3^/kg^64^. Cell lineages were visualized using a customized software based on Loon^81^.

### Engraftment Efficiency Assay

All protocols described in this and other sections regarding animal studies were approved by the UCSF Institutional Animal Care and Use Committee. Ethical endpoint for tumor transplantation experiments was reached when a tumor was 2.5 cm or more in any single dimension. Human melanoma cell lines with fluorescent BRN2 reporter were further transduced with pHIV-Luc-ZsGreen (Addgene 39196) and selected with neomycin (800 μg/mL) for 3 weeks. GFP high expressing cells were isolated by FACS and expanded. mCherry reporter high and reporter low cells were then FACS isolated as described above. 2 million cells from each group were injected into the flank of adult male NOD *scid* gamma mice at a site distant from the lungs. Mice were imaged in a PerkinElmer IVIS Spectrum Imaging System for luminescence every 2 weeks, 15 minutes after subcutaneous injection of 100 μl (150 mg Luciferin/kg body weight) of D-luciferin (Goldbio LUCK-1G). After the final week, mice were immediately sacrificed and organs were harvested and imaged. All animal studies were conducted by the UCSF Preclinical Therapeutics Core, of which the technicians were blinded to the cell types and hypotheses. Each group initially contained seven mice.

### Single Cell RNA Sequencing

scRNASeq libraries were prepared from freshly disassociated cell cultures according the BD Rhapsody System mRNA Whole Transcriptome Analysis manufacturer’s protocol (selected parameters: 11 cycles for sample tag PCR1 and 13 cycles for RPE PCR) and sequenced on an S4 Flow Cell of a NovaSeq6000. Raw data were processed using the BD Rhapsody WTA Analysis pipeline (https://igor.sbgenomics.com/public/apps/jiewho/bd-public-project/bd-rhapsody-wta-analysis-pipeline) which generates BAM files and single count matrices from raw FASTQ files. The BD Rhapsody output files (RSEC_MolsPerCell and Sample_Tag_Calls) were loaded into the Seurat 4.2.0 package in R^82^. Cells from 624Mel mcherry low and high were normalized using the sctransform v2 method^83^ and clustered using 15 dimensions and a 2.0 resolution with UMAP. Differentially expressed genes were identified using the default Wilcoxon Rank Sum test in Seurat. A gene set signature score was calculated for each cell by dividing the number of expressed genes in the set (with 1 or more read) by the total number of genes in the set. Module scores were added using ‘AddModuleScor’ in Seurat to identity gene sets with higher or lower expression than a random set of 100 genes^16^. Cell cycle scores were calculated using ‘CellCycleScoring’ in Seurat with the 2019 update of cell cycle genes. The cell annotations and count files from Jerby-Arnon et al^66^ were downloaded from GSE115978 at NCBI GEO and then combined into a Seurat object. The counts were normalized using the global-scaling normalization and module scores were added using AddModuleScore. The tSNE coordinates from the Single-cell portal (SCP109 Melanoma immunotherapy resistance) were added to the object for plotting module scores.

### Data Availability

The mRNA-sequencing data produced in this manuscript is available from GEO (GSE150582 and GSE230574).

### Statistical Analyses

For differential gene expression, p values were calculated with the DESeq2 (v1.30.1) default Wald test adjusted by the Benjamini-Hochberg method using a 5% false discovery rate (FDR)^84^. Pathway analyses were analyzed using the fast gene set enrichment package in R with a 10% FDR^85^. Gene set enrichment analyses were conducted using GSEA (v4.0.3, Broad Institute) Preranked tool with 1000 permutations. For other experiments, unpaired t-tests were used for Gaussian distributions and Mann-Whitney tests for non-Gaussian distributions or for samples of unequal variance. Exact p values were calculated as reported by Prism 8 (Graphpad).

## Acknowledgments

We utilized the Shared Resources for Research Informatics, High-Throughput Genomics and Bioinformatics Analysis, Preclinical Research, and Flow Cytometry and received direct financial support for the research reported in this publication provided by the Cell Response and Regulation Program at Huntsman Cancer Institute at the University of Utah, each supported by the National Cancer Institute of the National Institutes of Health under Award Number P30CA042014. The content is solely the responsibility of the authors and does not necessarily represent the official views of the NIH or other funding agencies.

## Author contributions

Conceptualization: YZ, RLJT; Methodology: YZ, RLB, TZ; Software: DL; Validation: MU; Formal analysis: RLB, MMKH, MH, TEM, FVR, MC, BKL, CS, XZ, TZ, RLJT; Investigation: YZ, RLB, MU, MMKH, TL, XZ, RLJT; Resources: RLJT; Data Curation: RLB, MC, CS, RLJT; Writing – Original Draft: RLJT; Writing – Review & Editing: YZ, RLB, MU, MMKH, FVR, RK; Visualization: RLB, MU, FVR, CS, TZ, RLJT; Supervision: AL, LSW, RK, RLJT; Project administration: RLJT; Funding acquisition: YZ, MWV, DG, RLJT

## Data and materials availability

All data are available in the main text or the supplementary materials. The mRNA-sequencing data produced in this manuscript is available from GEO (GSE150582 and GSE230574).

## Supplementary Materials

**Supplemental Figure S1: Flow analysis of clonally expanded and serially cultured reporter cells.** Flow analysis of clonally expanded then serially cultured SK-MEL-28 and WM793 cells with the BRN2 reporter.

**Supplemental Figure S2: Generation of horizon charts.** Graphical depiction of how the horizon charts portrayed in Fig. 3C are generated.

**Supplemental Figure S3: Quality control metrics for single cell sequencing.**

A) Percent of mitochondrial transcripts (pct_MT) and number of unique genes (nFeature_RNA) for single cell sequencing data. B) Representation of High^BRN2^ and Low^BRN2^ cells in each Seurat cluster. C-D) UMAP representations of signature scores for BRN2, AXL, MITF, and SOX10 regulated genes. E) UMAP with each cell colored by predicted cell cycle phase. F) The percent of cells in each Seurat cluster predicted in cell cycle phase.

**Supplemental Videos S1-2: Time-lapse imaging capturing bidirectional phenotype switching.**

**Supplemental Table S1: Differential expression analysis from bulk sequencing of mCherry high and low 624Mel cells.**

**Supplemental Table S2: Differential expression analysis from single cell sequencing of mCherry high and low 624Mel cells.**

**Supplemental Table S3: STR analysis of cell lines used in study.**

**Supplemental Table S4: Reagents used in study.**

