## Supplemental Figures 1-2 for "Bidirectional interconversion between mutually exclusive tumorigenic and drug-tolerant melanoma cell phenotypes"

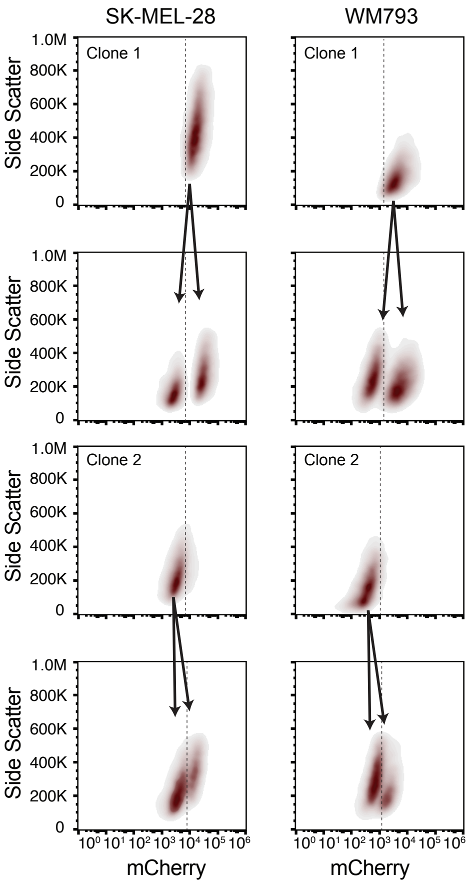


**Supplemental Figure S1: Flow analysis of clonally expanded then serially cultured reporter cells.**

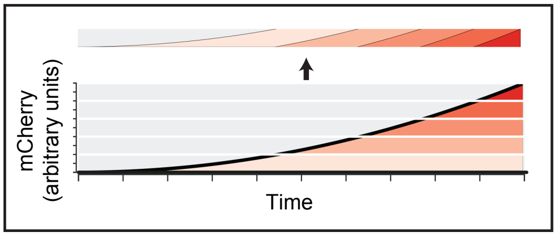


**Supplemental Figure S2: Generation of horizon charts**

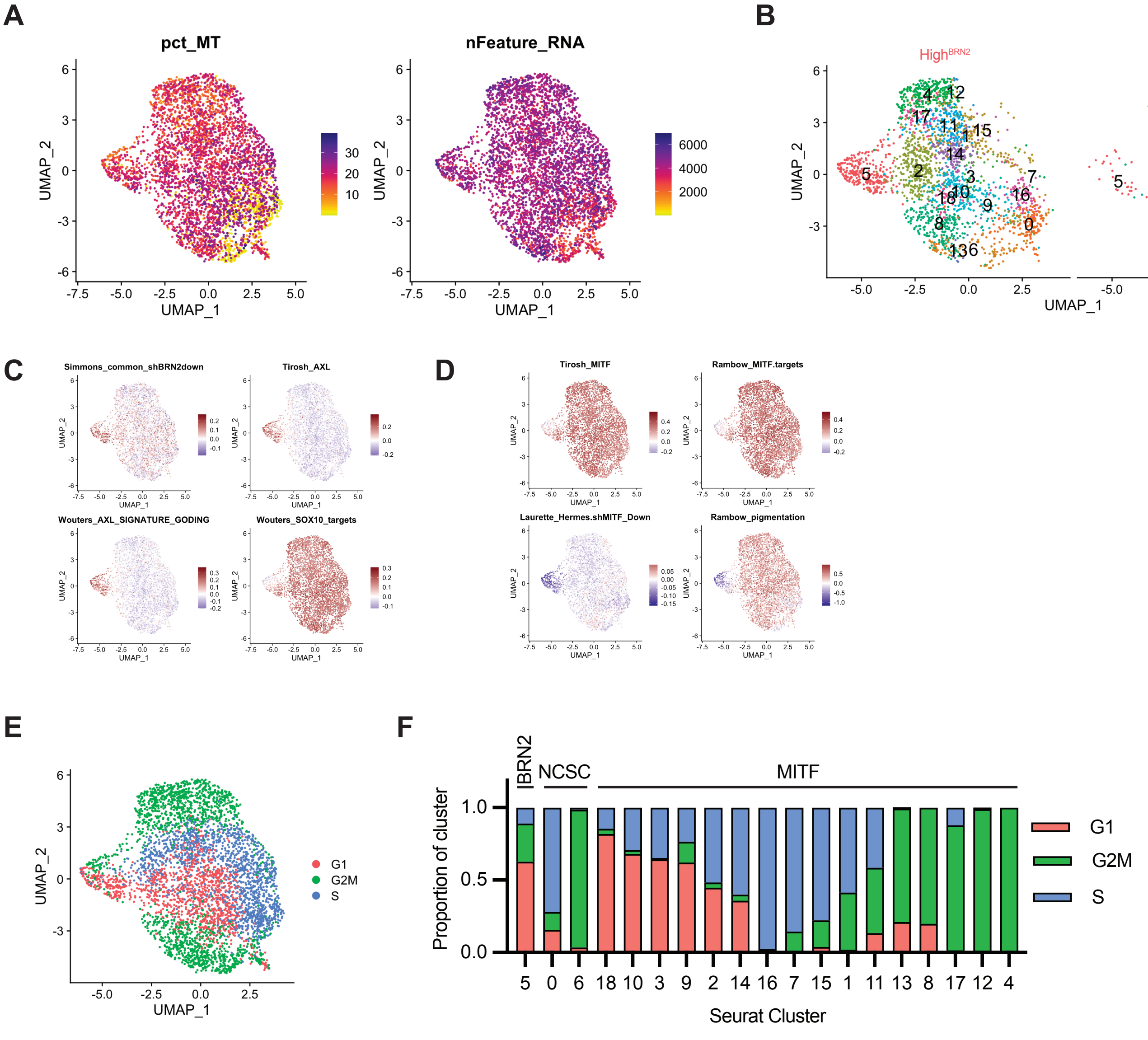


**Supplemental Figure S3: Quality control and characterization of single cell sequencing.**
